# Mother cells can establish slow growing lineages in clonal populations since their earliest division cycles

**DOI:** 10.1101/2023.04.20.537656

**Authors:** Irene Delgado-Román, María José García-Marcelo, Carmen Ruger-Herreros, Lidia Delgado-Ramos, Abhyudai Singh, Sebastián Chávez, Mari Cruz Muñoz-Centeno

## Abstract

Clonal populations exhibit phenotypic variation despite being composed of genetically identical cells under the same environmental conditions. Proliferation rate also shows this heterogeneity, but the underlying mechanisms remain poorly understood. In this study, we combined single-cell microencapsulation with confocal microscopy to develop a new experimental approach for analyzing budding yeast cell lineages and determining the age of every cell within each microcolony. We found that most slow-growing lineages are founded by young mother cells that have undergone only a few cell divisions, typically between 1 and 4. This reduction in proliferative capacity is linked to the expression levels of the cell cycle regulator Whi5, which increase with the number of replication cycles, even since the earliest stages. We also found that the increased levels of Whi5 are due to the higher accumulation of its mRNA during the S/G2/M phases of young mother cells compared to newborn cells. Our results show that the proliferative structure of a cell population is progressively shaped in each mitotic cycle, starting from the very first division, when a mother cell has the opportunity to establish a slow proliferating lineage. Possible mechanisms of Whi5 action to mediate this effect are discussed.

## Introduction

Heterogeneity can be defined as the presence of different characteristics within a population ^1–3^. Variability among cells within the same clonal culture can arise due to mutations accumulated during population growth. In addition to this genetic heterogeneity, which is relatively well understood ^4–9^, there is also phenotypical heterogeneity, observed in isogenic populations, which does not depend on mutations or environmental changes. In this latter case, heterogeneity can manifest in various aspects of the cell phenotype, such as shape, size, or differential expression of specific genes ^10–13^. Phenotypic heterogeneity can also influence the ability of isogenic cells to divide and proliferate.

Proliferative heterogeneity explains variations in growth rate within a clonal cell culture, as repeatedly observed in the budding yeast *Saccharomyces cerevisiae* ^12,14^. Previously, we described how the encapsulation of single cells from a yeast clonal population in alginate microspheres results in a wide range of microcolony sizes due to differences in proliferation rates^15^.

The first events leading to these different proliferation patterns may arise from small stochastic fluctuations in the expression profile, commonly referred to as “noise,” which result in differences in proliferation^10,16^. If these fluctuations affect critical genes, as has been described for the histone acetyltransferase *SIR2*, epigenetic changes can lead to stable expression patterns that alter proliferation rates^17–19^.

Although the whole *S. cerevisiae* lifespan has been studied, with significant insights gained from mother cells after a high number of divisions, the impact of the earliest mitotic cycles (from the first to the fifth first divisions) on proliferation rates remains unexplored. Understanding these initial stages is crucial to deeply understand the proliferative heterogeneity within clonal exponential cultures, where the majority of cells are newborn or recent descendants, while older mother cells are vastly outnumbered. ^20^.

In previous works, we performed transcriptomic analyses of yeast microcolonies based on their size, revealing differential gene expression profiles between fast- and slow-growing clonal subpopulations ^15^. The most differentially overexpressed gene in small, slow-growing microcolonies was *WHI5*, a G1-S transition regulator that represses the expression of G1 cyclins by inhibiting the transcription factors SBF and MBF^21–23^, in a manner similar to how Retinoblastoma protein controls G1-S transition in mammalian cells^24,25^. It has been proposed that the levels of Whi5 acquired by daughter cells during budding are essential for size-dependent entry into G1 ^26^.

These results led us to explore whether the growth rates of microcolonies might be influenced by the age of the founder cell and whether variations in Whi5 levels might contribute to the proliferative heterogeneity of populations that are genetically identical but intrinsically diverse in replicative age. To address these questions, we combined single-cell microencapsulation with confocal microscopy and found that slow-growing microcolonies are frequently founded by mother cells rather than the more abundant newborn cells. Surprisingly, we also found that the founders of slow-growing microcolonies were not at all aged cells but rather young mother cells with just 1 to 4 division cycles. Furthermore, we discovered that the reduced proliferative capacity of microcolonies founded by young mother cells is linked to Whi5 expression levels, which increase with mother cell age from the very first division due to the higher accumulation of its mRNA after the S phase. Our results demonstrate that the proliferative structure of a cell population is shaped during every mitotic cycle, beginning with the very first divisions, when a mother cell has the opportunity to increase Whi5 expression levels and establish a slow-proliferating lineage. Thus, proliferative heterogeneity arises from a programmed system that introduces diversity in the expression levels of a cell cycle regulator from the very first round of division.

## Results

### A new method for analyzing the replicative age of cells within a yeast microcolony: establishing growth patterns

We have already shown that encapsulated microcolonies can be handled like particles suspended in a fluid, allowing, for example, their analysis in a large particle flow cytometer ^14^. We conceived that this would also enable the 3D analysis of individual microcolonies by confocal microscopy. We wondered whether such a 3D analysis would help us explain why some cells generate microcolonies at a lower proliferation rate than others.

We first established the criteria to microscopically detect those microcolonies with the most reduced proliferative rate, which generate smaller microcolonies. As described in Supplementary figure S1A, we considered “small” (S) microcolonies those with a diameter lower than 25 µm after encapsulation and 13 hours of incubation in optimal conditions. We considered “big” (B) microcolonies those larger than 50 µm in diameter, and “medium” (M) microcolonies those in between 25 and 50 µm in diameter. As shown in Supplementary figure S1B, nearly 30% of the microcolonies generated fell into the small category, exhibiting low proliferative capacity; more than 50% met the size criteria for the medium category, and less than 20% were classified as big.

In order to gain insight into the genealogical structure of the microcolonies, it is important to determine the replicative age of their cells. The number of bud scars serves as an excellent marker of replicative age. We demonstrated that the bud scars of encapsulated cells can be optimally detected after calcofluor staining, without compromising microcolony integrity, as shown in Supplementary figure S2.

After calcofluor staining, small microcolonies were analyzed under a confocal microscope, as shown in Figure 1A. For each small microcolony, we counted the total number of cells and bud scars, as well as the number of bud scars present in each cell by examining all Z-axis images. We collected this data for more than 100 small microcolonies and analyzed the distribution of bud scars per cell to infer genealogical patterns (Supplementary Table S1 and supplementary data: all images used in this study are available at http://www.ebi.ac.uk/biostudies/ with accession number S-BSST1071). As a quality control, we calculated the total number of bud scars (n) and verified whether it was consistent with the total number of cells counted (N). The number of scars should always be greater than the number of cells minus two (n > N -2). Microcolonies that did not meet this condition, likely founded by aggregated cells, were discarded.

**Figure 1.**
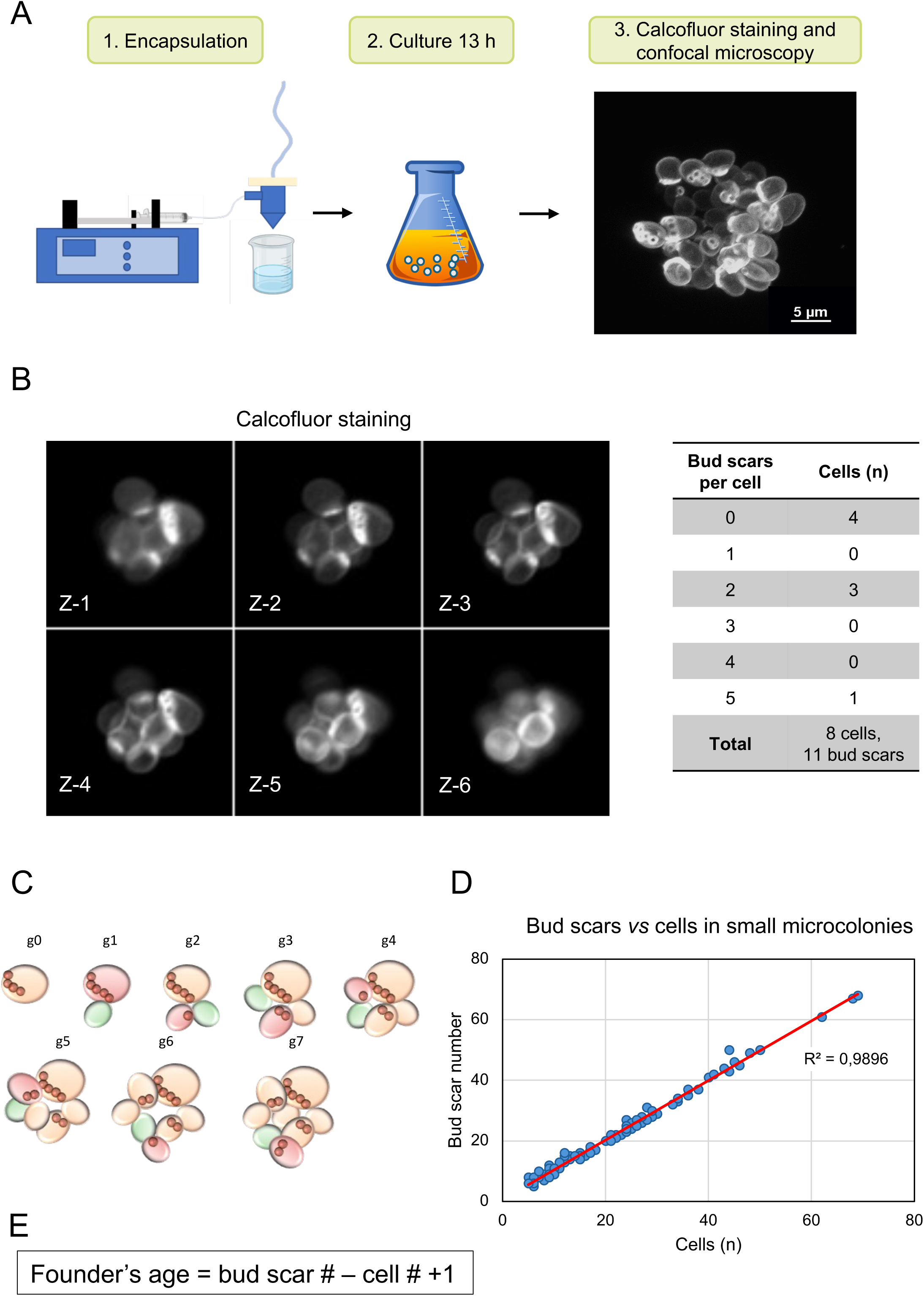
Accurate determination of the replicative age of the founder cell and the genealogy of the microcolony. (A) Schematic representation of the microcolony analysis methodology. The microencapsulation conditions were set to ensure that a single cell from the clonal cultures was present in each capsule. The capsules were then collected and cultured together in an appropriate medium for 13 hours at 30 °C. Finally, fixed microcolonies were filtered, stained with calcofluor, and analysed using confocal microscopy. (B) The left panel shows confocal images (Z-stacks Z1– Z6) of a small microcolony taken after 13 hours of growth and stained with calcofluor to facilitate bud scar counting. This microcolony was founded by a non-newborn cell at generation 0 (g0). The right panel shows an overview of the counting methodology used to identify the founder cell and its genealogy for the same microcolony. For each microcolony, the total number of cells and bud scars is shown, as well as the number of cells ranked by bud scar number. (C) Graphical representation of the development of the microcolony shown in (B) across generations (g0–g7). Each generation represents a cell division event. The new daughter cell generated in each mitotic event is depicted in green, and the corresponding mother cell in red. Bud scars are represented in brown. In this case, no division restrictions were detected among the microcolony cells. We refer to this non-restricted proliferation pattern as isotropic. (D) Plot showing the total number of cells versus the total number of bud scars for each microcolony analysed. (E) Formula used to estimate the replicative age of the microcolony founder cell at the time of encapsulation as a single cell.

As mentioned above, for each small microcolony, we counted the total number of cells and bud scars, as well as the number of bud scars present on each cell. An example is shown in Figure 1B. These data allow us to reconstruct the microcolony’s genealogy. In most cases, the distribution of scar numbers was consistent with classical exponential growth. In these cases, unbudded newborn cells (with no scars) were the most frequent category within this microcolony, but never accounted for more than 50%. These microcolonies also contained decreasing proportions of cells with one or more buds. The cell with the highest number of buds was identified as the likely founder of the colony. We refer to this pattern of growth as “isotropic,” where every cell within the microcolony contributes proportionally to its formation because all cells divide at the same rate. The confocal analysis of a representative example of an isotropic small microcolony is shown in Figure 1B. In this example, the small microcolony is composed of 8 cells, indicating that after the microencapsulation of a single cell, 7 divisions took place. The cell exhibiting 5 bud scars was identified as the founder. Four cells were newborns with no bud scars, and the other 3 cells showed an intermediate number of bud scars, indicating that they had divided during the development of the microcolony. The potential sequence of division events that explains the distribution of scars is shown in Figure 1C. Under this isotropic pattern, although the proliferation rate was low and only produced 8 cells in 13 hours, no particular cell in the microcolony showed a stronger restriction to divide.

A second growth pattern, although a minority, was also observed in some small microcolonies. A representative example is shown in Figures 2A and 2B. In this case, the founder cell, after encapsulation, underwent several divisions, but its daughter cells either did not divide or showed severe restrictions in doing so. We have referred to this growth pattern as “anisotropic”. This kind of microcolonies was not the result of artefactual co-encapsulation of two aggregated cells with different age, an event with a much lower probability to take place (see Materials and Methods).

**Figure 2.**
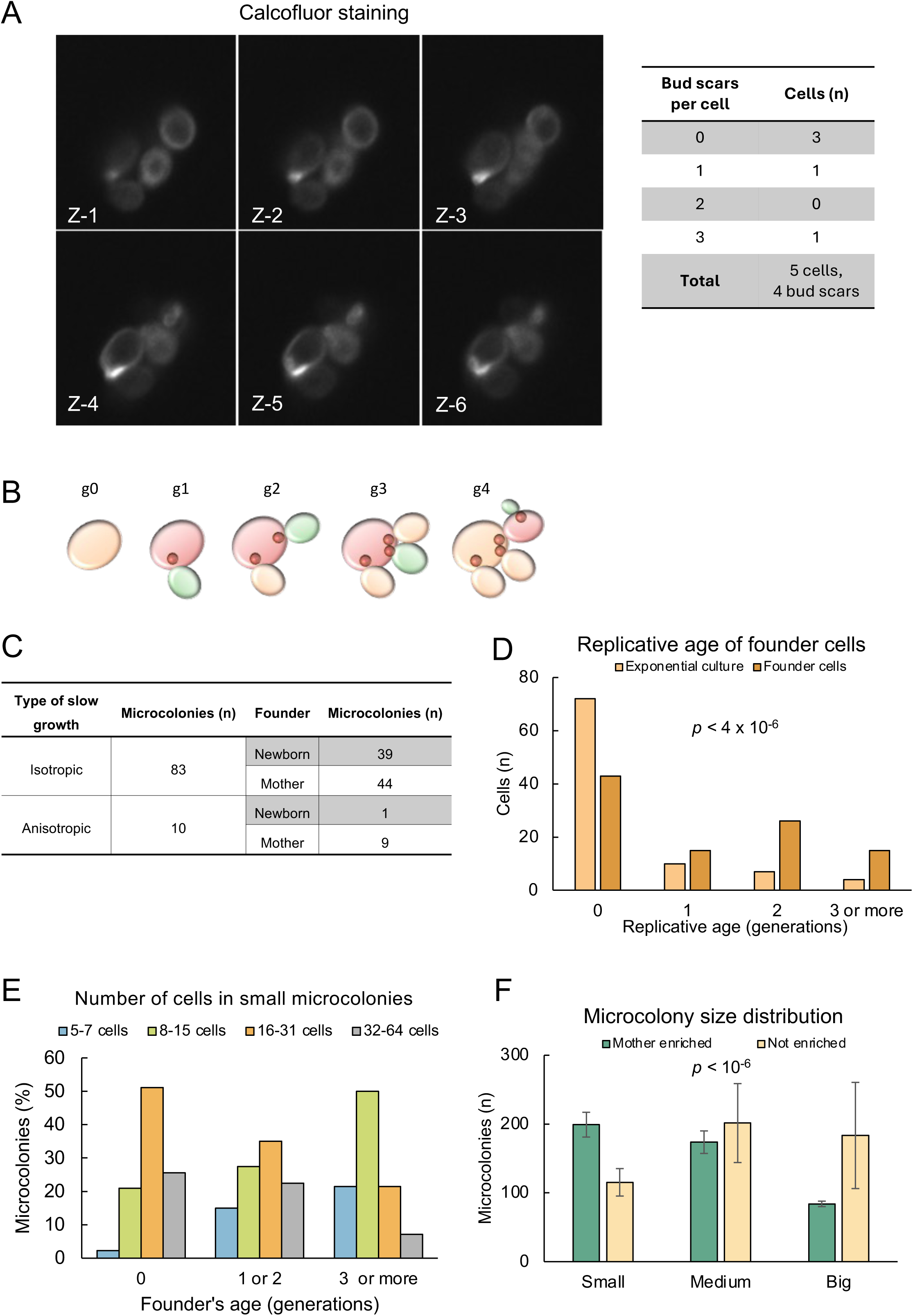
Small microcolonies are more frequently formed by young mother cells than by newborns. (A) The left panel shows confocal images of a small microcolony taken after 13 hours of growth and stained with calcofluor. This microcolony was founded by a newborn cell (g0). The right table shows the total number of cells and bud scars, as well as the number of cells ranked by bud scar number. (B) Graphical representation of the development of the microcolony shown in (A) across generations (g0–g4). Daughter cells generated in generations 2 and 3 did not divide further, indicating a proliferation restriction in this microcolony. This pattern of restricted proliferation is called anisotropic. (C) Growth patterns and founder cells of slow-proliferation microcolonies. Small microcolonies were classified as ‘isotropic’ when most cells within the microcolony divided equally. They were classified as ‘anisotropic’ when one cell (either the founder cell or a first-born cell) underwent most divisions within the microcolony. In all cases, the founder cell was identified and classified as either newborn or non-newborn (mother). Regardless of the growth pattern, slow-proliferation microcolonies tend to be founded by mother cells. See the text for details. (D) Distribution of founder cells (dark orange) according to their replicative age at the time of microcolony formation (n = 96), compared to the initial exponential culture prior to encapsulation (light orange) (n = 96). The two distributions exhibited a significant statistical difference according to a square-chi test (*p* = 3.6·10⁻⁶). (E) A comparison of the distributions of the replicative age of the founder cells at the time of founding among small microcolonies of different sizes. Small microcolonies (those with fewer than 65 cells after 13 hours of growth) were classified into four categories according to their number of cells: 32–64, 16–31, 8–15, and 5–7. The smaller the microcolony, the higher the proportion of non-newborn founder cells. There is a progressive increase in the proportion of microcolonies founded by mother cells with a higher number of replication cycles in the smaller categories. (F) Encapsulation of a population of cells enriched in young mothers with one or two division cycles following the Mother Enrichment Programme (see Supplementary Figures S4A–B) produced a higher proportion of small microcolonies. The distribution of microcolonies according to their size (small, medium or large; see Supplementary Figures S1A and S1B) is shown for a control population (untreated, light yellow) and a mother-enriched population (treated for 3 hours, green). Cultures enriched in young mothers produced a higher proportion of small microcolonies than the non-treated control. Remarkably, more than 60% of the cells in this mother-enriched culture had a replicative age of between one and five, meaning they were young mothers, not old cells. The two distributions exhibited a significant statistical difference according to a chi-square test (*p* < 10⁻⁶).

The confocal analysis and data from a representative example of an anisotropic small microcolony are presented in Figure 2A. In this example, the small microcolony contained 5 cells, indicating that 4 divisions occurred after the encapsulation of a single cell. The cell with 4 bud scars was identified as the founder. Three cells were newborns with no bud scars, and one additional cell had a single bud. The distribution of bud scars per cell indicates that all but one of these divisions occurred in the founder. A potential sequence of division events that explains the anisotropic development of this microcolony is depicted in Figure 2B.

Once we defined these two growth patterns, we quantified how many of the small microcolonies analyzed corresponded to each of them. We observed that most of the microcolonies (83 out of 93) exhibited an isotropic growth pattern, while only 10 displayed the anisotropic one (Figure 2C, Supplementary Table S1, and Supplementary images at http://www.ebi.ac.uk/biostudies/).

### Small microcolonies are more frequently founded by non-newborn cells

To determine whether replicative age, starting from the first divisions, contributes to the proliferation heterogeneity observed in clonal cells, we decided to identify the founder cell in each small microcolony and estimate its replicative age at the time of foundation. As mentioned above, the total number of bud scars per microcolony is always similar to or slightly higher than the number of cells (Figure 1D), with this difference depending on the replicative age of the founder cell. To estimate its replicative age at the time of encapsulation (A), we reasoned that it should correspond to the difference between the total number of bud scars in the microcolony (n) and the total number of cells (N) plus one, as illustrated in Figure 1E (A = n -N + 1) (see Materials and Methods).

In the example shown in Figures 1B-C, the 8 cells of the microcolony exhibited a total of 11 bud scars, indicating that the founder cell had already divided three times before founding the microcolony. In the second example (Figures 2A-B), the 5 cells showed 4 bud scars, suggesting that the founder cell was a newborn. Using this simple calculation, we were able to estimate the replicative age of the founder cells in the 96 small microcolonies analyzed (Supplementary Table S1). According to our analysis, most small microcolonies were founded by non-newborn cells. This result was observed in both isotropic (more than half) and anisotropic (nine out of ten) microcolonies (Figure 2C).

To really understand the relevance of this proportion of non-newborn cells as small microcolonies founders, we compared the age distribution of the founder cells (at the time of microencapsulation) to that of cells from the same exponential liquid culture from which the encapsulated cells were taken. The results are presented in Figure 2D. As previously described ^27^, most of the cells in the exponential culture were newborns (about 64%) and decreasing proportions of cells containing from 1 to 3 and more bud scars were also detected (Figure 2D in light orange). Since we used exponential liquid cultures for cell microencapsulation, we would expect the same age distribution in the founder cells of small microcolonies. In contrast, the age distribution of the small microcolony founder cells was significantly different (p < 3.6 x 10⁻⁵) (Figure 2D in dark orange): there were much fewer newborn founders than in the liquid culture, and a higher frequency of founders that had undergone some divisions before encapsulation than expected (Figure 2D and Supplementary Figure S3).

To reinforce this conclusion, we also studied in more detail the size of these small microcolonies in relation to the age of the founder cell, and the results are shown in Figure 2E. We found that most small microcolonies founded by newborn cells consisted of more than 16 cells, whereas most of those founded by mothers with three or more previous divisions exhibited fewer than 16 cells. Small microcolonies founded by young mothers (with one or two previous divisions) showed an intermediate size distribution. These results indicate that a slight increase—such as one or two divisions—in the founder cell is associated with a decrease in the proliferation rate of the microcolony. Additionally, these findings suggest that the final size of the microcolony is influenced by the number of generations of the founder cell at the moment of encapsulation. Thus, a single division of a newborn cell is sufficient to enhance the likelihood of continued proliferation at a reduced growth rate, ultimately leading to the formation of smaller microcolonies.

The fact that microcolonies founded by very young mother cells exhibited a significantly higher tendency to be small suggests a negative influence of replicative age on cell proliferation from the earliest stages of the lifespan. To confirm this first conclusion of our study, we utilized the “Mother Enrichment Program” (MEP), an inducible genetic system regulated by the presence of estradiol. This system restricts the replicative capacity of daughter cells while allowing mother cells to maintain a normal replicative lifespan (Supplementary figure S4A) ^28^. In the absence of estradiol, both mother and daughter cells can divide normally. However, in the presence of estradiol, daughter cells become unviable, and the population becomes enriched in mother cells of increasing age.

We hypothesized that a slightly aged population, not dominated by newborn cells but rather by young mother cells that had undergone one or two divisions, would result in an increased proportion of slow-proliferating microcolonies after encapsulation. First, we optimized the estradiol treatment required for the MEP strain DNY51 to achieve the mild aging effect we needed. We found that 3 hours of incubation with estradiol was optimal for generating a population with half the number of newborn cells seen in the control culture, along with a significant enrichment of mother cells with one or two bud scars (Supplementary Figure S4B). While more than 60% of cells in the untreated population were newborn, in this estradiol-treated culture, 60% of the cells had undergone at least one budding cycle (Supplementary Figure S4B).

These two cell populations were encapsulated and incubated for an additional 13 hours in the absence of estradiol to allow microcolony formation without further selection against daughter cells. Microcolonies were then classified according to their size, as described in Supplementary Figure S1A, and compared (Figure 2F). We found that the proportion of small microcolonies increased in the particles seeded with estradiol-treated cells (in green) compared to the control samples (in light yellow), becoming the predominant microcolony category in this condition. The overall distribution of microcolony size was significantly altered by the limited number of division cycles previously undergone by the founder cells (Chi-square test, *p* < 0.001). This result strengthens the link between replicative age and proliferation rate, even at the very early stages of a mother cell’s lifespan, indicating that, once a daughter cell divides, there is a significant probability of establishing a lineage with a low proliferation rate across generations.

### Whi5 levels set the proliferation rates of microcolonies

As mentioned in the Introduction, in our transcriptomic analysis we found that slow- and fast-proliferating microcolonies exhibited differential transcriptomic signatures. Specifically, *WHI5* was the most overexpressed gene in small slow-proliferating microcolonies^14^. In budding yeast, Whi5 is a G1/S transition inhibitor ^25^ that contributes to both the determination of critical cell size at START and cell fate ^29–31^. We decided to confirm that the correlation between Whi5 high levels and slow-proliferating (small) microcolonies was causative, indicating that Whi5 levels determined slow proliferation rather than the other way around. To do this, we used the *S. cerevisiae* KSY098 strain^32^, where *WHI5* gene expression is regulated by a chimeric regulator containing an estrogen-binding domain and responds to estradiol in a dose-dependent manner^33^ (Figure 3A).

**Figure 3.**
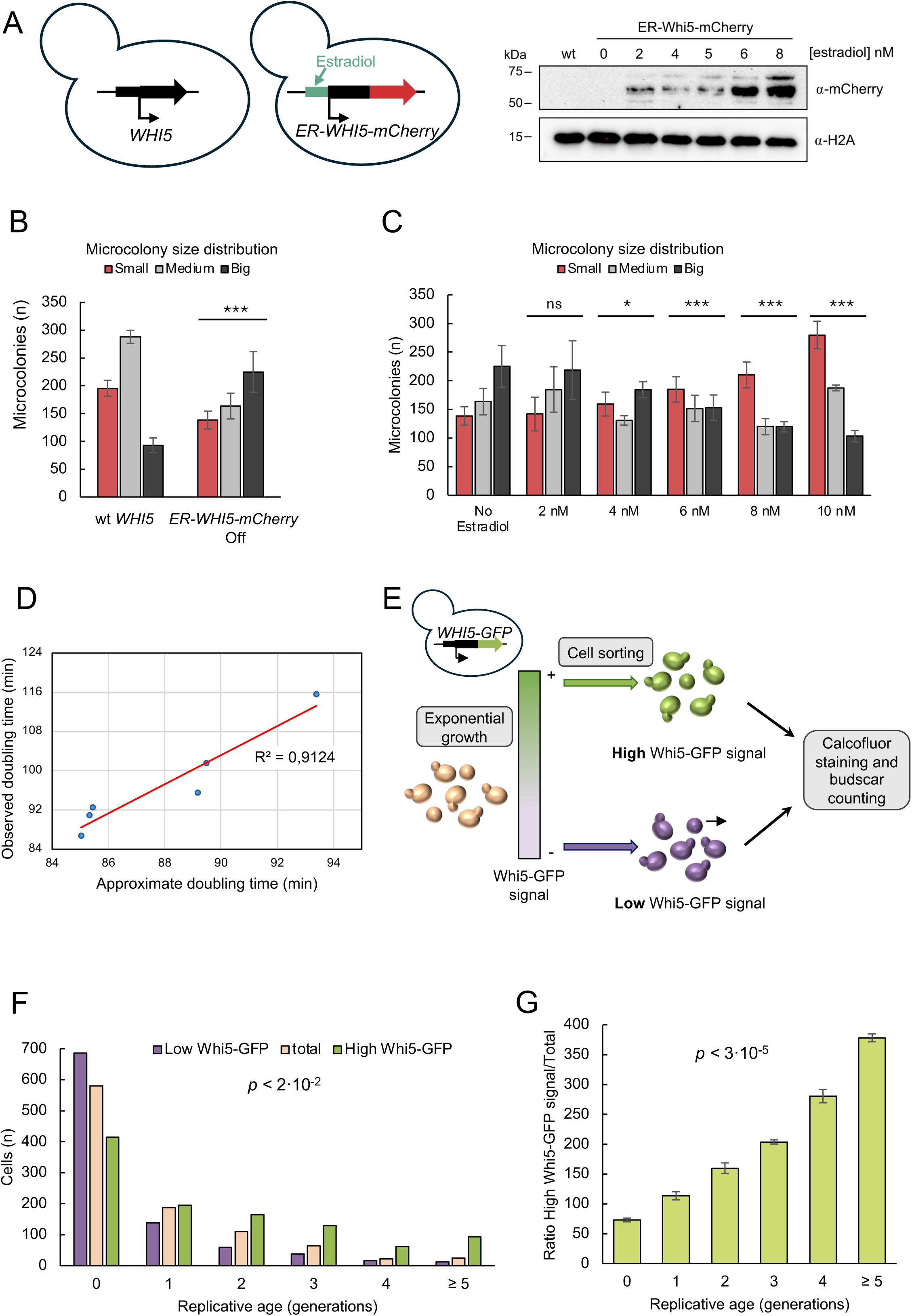
Upregulation of Whi5 levels results in a higher frequency of slow-proliferating microcolonies founded by non-newborn cells. (A) Left panel shows a schematic representation of the Whi5 region in the KSY098 strain. The endogenous locus, including the promoter, has been replaced by a system that controls Whi5 expression using estradiol and adds the mCherry epitope to the C-terminal region (see Table S3). Right panel shows a Western blot of the inducible Whi5 strain (KSY098) at different concentrations of estradiol. Whi5 accumulates in a dose-dependent manner according to the increasing concentration of estradiol in the medium. (B) A comparison of the size distributions of microcolonies formed by KSY098 cells grown without estradiol and a wild-type (wt) control. Compared to the wt profile, the lack of Whi5 expression produced a significantly different pattern characterised by an increase in large microcolonies (black) and a decrease in small microcolonies (red). Chi-square tests was performed to determine statistical significance (*p-*value lower than 5·10⁻⁴). (C) Microcolony size distribution profile of the KSY098 strain cultured without estradiol (control) or with 2., 4, 6, 8 or 10 nM estradiol, respectively. As the concentration of estradiol increases and the expression levels of Whi5 increase proportionally, a progressive increase in the proportion of small microcolonies is detected. Chi-square tests were performed to determine statistical significance. *, **, *** and ***** indicate *p-*values lower than 5·10⁻², 5·10⁻³, 5·10⁻⁴ and 5·10⁻⁶, respectively. (D) Correlation between measured doubling times of KSY098 cultures treated with the indicated estradiol concentrations and the estimated doubling times from mathematical modelling of results shown in C (see text). (E) Experimental procedure for cell sorting of Whi5-GFP according to the GFP intensity signal. An exponential Whi5-GFP culture in YPD was collected and sorted according to its GFP signal. Three samples were collected: cells with the highest Whi5-GFP expression; cells with the lowest Whi5-GFP expression; and a sample of the entire population to serve as a control for the culture (total). Each sample was then stained with calcofluor and separately analysed. (F) The replicative age distribution, represented by the number of bud scars, for the entire cell population (peach) and the Whi5 low-expressing (purple) and high-expressing (green) subpopulations. Bud scars were counted in three independent replicates of the experiment. Approximately 1000 cells from each category were counted in total. The subpopulation of cells expressing higher levels of Whi5-GFP was enriched in young mothers with two or more division cycles compared to the subpopulation expressing lower levels of Whi5-GFP, which was enriched in newborns. The difference between these two categories in terms of replicative age was statistically significant according to a chi-square test (*p* lower than 0.02). (G) The ratio of cells with high Whi5 expression relative to the whole population in each replicative age group. For each biological replicate, the proportion of cells of a certain age in the high Whi5-GFP sample was divided by the corresponding proportion in the whole population. The average and standard deviations are shown. The frequency of high Whi5 expression increases with the number of mitotic cycles undergone since the first division. The statistical significance of the overall differences was confirmed with an ANOVA test (*p* lower than 2.6 x 10⁻⁵).

First, as shown in Figure 3B, we compared the microcolonies produced by encapsulated KSY098 cells cultured in the absence of estradiol (referred to as *WHI5* Off), which therefore express undetectable levels of Whi5 (Figure 3A, right panel, line two), to those produced by isogenic W303 cells, which express *WHI5* from its own promoter (referred to as WT-*WHI5*). The lack of Whi5 expression resulted in a significant shift in the microcolony distribution towards the big category, at the expense of the medium and small ones (Figure 3B). This result confirmed that Whi5 plays a role in the proliferation heterogeneity detected in our microcolony assay and that its expression is detrimental to the fast-growing subpopulation.

This result prompted us to hypothesize that, on the contrary, higher levels of Whi5 should favor slow-proliferating or small microcolonies. To test this hypothesis, we encapsulated exponentially growing KSY098 cells in the absence of estradiol (and thus not expressing *WHI5*). Single-cell seeded particles were then incubated either without estradiol or with estradiol concentrations ranging from 2 to 10 nM (see Supplementary Figure S5 panel A for a synopsis of the experiment). Interestingly, cells incubated with 4 nM estradiol after encapsulation showed a significant shift in microcolony distribution toward the small category (slow-proliferating rate), at the expense of the large category (Figure 3C). Higher concentrations of estradiol, which resulted in stronger expression of Whi5 (Figure 3A), also caused an increase in the frequency of small microcolonies (Figure 3C).

Cell volume was also measured at the different estradiol concentrations. As expected, cells did not undergo any volume reduction with Whi5 levels, but a moderate increase, supporting the notion that small microcolonies arise from lower number of cells due to slower proliferation of the lineage, and not from the same number of smaller cells (Supplementary Figure S5B left panel). In fact, observation under the microscope of microcolonies of different sizes produced with the same estradiol concentration did not reveal streaking differences in cell size (Supplementary Figure S5B right panel). Notably, cytometry profiles of cultures with increasing Whi5 levels were consistent with the modest volume increase observed: even at the highest Whi5 levels, cells continued to cycle and the expected slight increase in G1 was observed (Supplementary Figure SC).

This increase in the frequency of slow-proliferating lineages caused by moderate Whi5 upregulation was expected to affect the overall growth rate of the cell population. Following mathematical modelling, we estimated the overall growth rates that small/medium/big microcolony distribution would predict in each of the experimental conditions shown in Fig 3C. The resulting expected doubling times exhibited high correlation with doubling times measured from exponential liquid cultures (Figure 3D). Predicted doubling times, however, were lower than observed, suggesting additional limitations for growth that were not considered in our mathematical model.

### The proportion of cells expressing high levels of Whi5 increases after the first mitotic cycles

The results described above confirmed that small microcolonies are associated with high Whi5 levels and that this correlation is causative. We have also shown in this work that such small slow-proliferating microcolonies were frequently founded by cells that had undergone more than one previous division cycle. We wondered whether these two phenomena were linked in such a way that Whi5 levels increase with the number of mitotic cycles performed by a newborn cell.

To test this hypothesis, we analyzed cells expressing a Whi5-GFP fusion as the sole source of Whi5, controlled by the endogenous *WHI5* promoter. Exponentially growing Whi5-GFP cells were sorted and collected based on their GFP signal, as described in detail in the Materials and Methods section (see Supplementary Figure S6 and Figure 3E for a summary of the experiment). We collected three different types of samples: i) cells with the highest Whi5-GFP expression (represented in green); ii) cells with the lowest Whi5-GFP expression (represented in purple); iii) a random sample of the entire population as a control (represented in peach) (Figure 3E-F and Supplementary Figure S6). Each sample was then stained with calcofluor and analyzed under the microscope to determine the replicative age distribution of cells. We found the expected distribution of bud scar counts in the control sample (peach bars in Figure 3F). In contrast, the sample with low Whi5-GFP levels showed a significant enrichment of newborn cells (purple bars in Figure 3F). Furthermore, the replicative age distribution of cells sorted for high Whi5-GFP levels displayed a clear reduction in newborn cells and an enrichment of cells with two or more division cycles (green bars in Figure 3F). To analyze these results in more detail, Figure 3G shows, for each replicative age category, the ratio of cells with higher Whi5 levels (green) versus the total number of cells in that category. Taken together, these results suggest that mother cells increase Whi5 expression as they undergo division cycles, starting from the earliest stages of their lifespan.

### Young mother cells overexpress Whi5 before cytokinesis

The correlation observed between decreased proliferation rate and higher Whi5 levels in young mother cells prompted us to investigate how this increase in Whi5 occurs. It is well-established that *WHI5* mRNA accumulates during the S-G2-M phases of the cell cycle ^34,35^. Thus, we decided to examine whether the duration of these cell cycle phases changes when a newborn cell undergoes its first rounds of division.

First, we investigated potential differences in the cell cycle phase profile in wild-type cells during the earliest replicative age stages. After calcofluor staining, cells were analyzed for cell phase morphology, and bud scars were counted (Figure 4A). As previously described, newborn cells are enriched in G1, as they need to reach the size threshold necessary for cell cycle progression. Consequently, the proportion of newborns in G1 (more than 60%) was much higher than that of mothers with one division (less than 25%) (Figure 4A). However, the G1 proportion increased again in mother cells with two divisions, up to more than 40% after three division cycles (Figure 4A). It is important to emphasize that the data presented in Figure 4A represent the relative duration of G1 compared to S/G2/M, rather than absolute time values, which must be estimated using time-lapse strategies. Analyzing published data obtained with this approach, we found significant alterations in cell cycle progression across early replicative ages (0–7 divisions). The analyses revealed marked changes in the average duration of S phase and increased heterogeneity in G2/M as replicative age advanced^36^.

**Figure 4:**
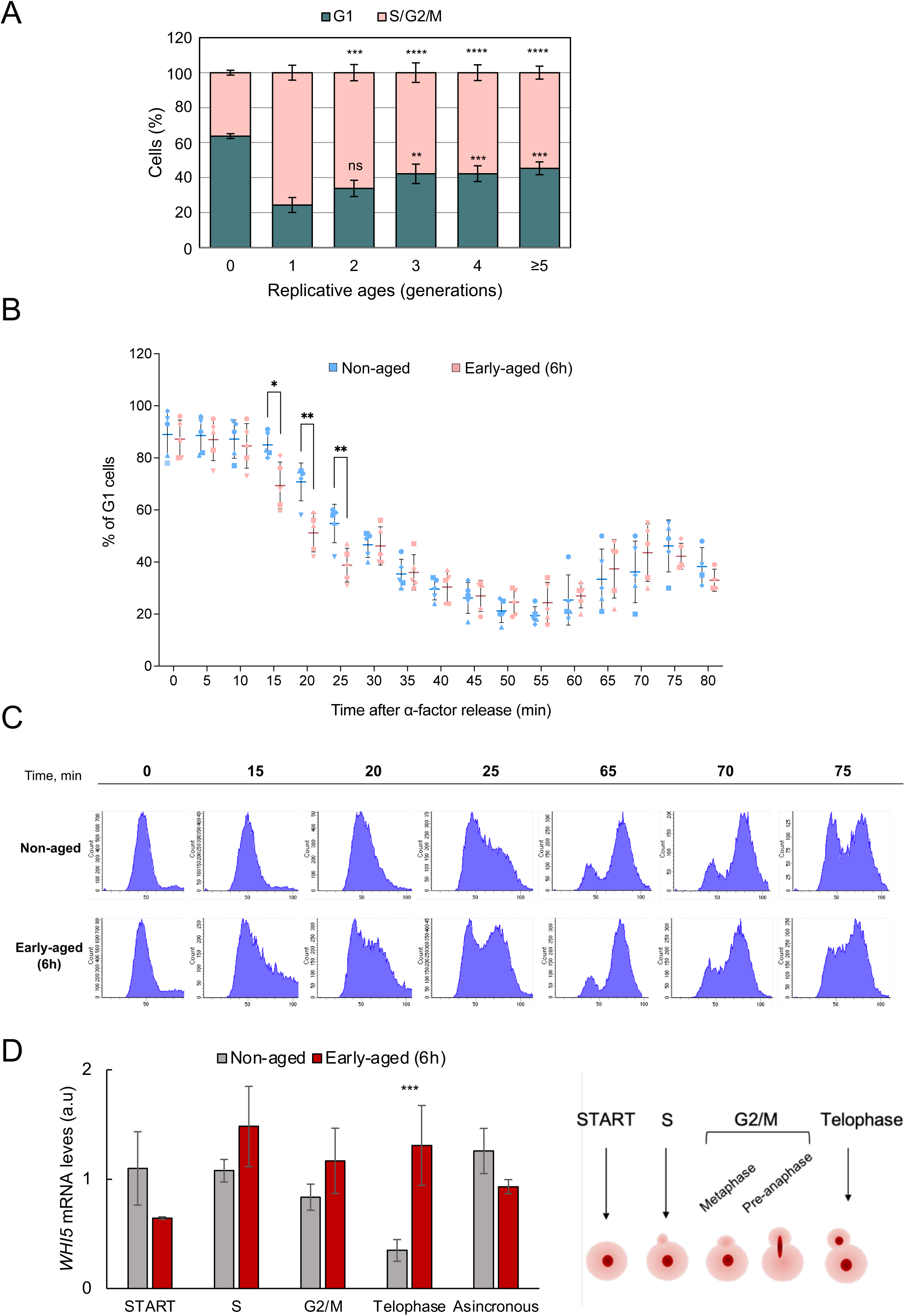
Young mother cells spend a longer time in the S, G2 and M phases of the cell cycle than newborn cells and accumulate higher levels of *WHI5* mRNA before cytokinesis. (A) Cell cycle profile of a wild type according to the replicative age of the cell. The cells were first stained with calcofluor to quantify their bud scars and then categorised by cell cycle phase based on cell and bud morphology and size analysis. One-way ANOVA test were performed with the absolute cell numbers to determine statistical significance between G1 or S/G2/M phases at different replicative ages. **, *** and **** indicate *p*-values of less than 1·10⁻², 1·10⁻³ and 1·10⁻⁴ respectively. (B) Cell cycle progression through S/G2/M of cultures enriched in young mother cells with three to five divisions using the biotin/streptavidin method (pink), compared to control populations dominated by newborns (blue) (see Supplementary Figure S7). Cells released from alpha-factor synchronisation in START were sampled at the indicated times and analysed by flow cytometry. The proportion of cells in G1 of five independent replicates is shown, with each replicate represented by a different symbol. For the first 10 minutes, both types of culture remained synchronised in G1; however, from minute 15 onwards, mother-enriched cultures consistently entered the S phase, whereas control cultures did not until after minute 20. After 35 minutes, cells from both cultures progressed through G2-M in an almost identical manner until the next G1 phase (after 65 minutes). (C) Flow cytometry of critical time points detected in B. See also Supplementary Figure S8B. (D) Relative *WHI5* mRNA levels analysed by RT-qPCR in early-aged cultures (mother-enriched) (red) and control cultures (grey). Samples from three replicates of each type of culture, manipulated as in A, were taken at the times described in Table S2 and analysed by flow cytometry and microscopy to ensure they corresponded to the exact step of the cell cycle being examined. We observed that *WHI5* mRNA peaked in the S phase in control cultures, but remained high until telophase in mother-enriched cultures.

In order to test whether the relative variation in the G1/G2-M proportion in young mothers reflected an absolute change in the duration of the cell cycle after START, we performed a G1 synchronization and release assay. To achieve this, we first carried out the previously described biotin-streptavidin purification assay (see Materials and Methods) to enrich cultures with the aforementioned very young mothers. Differences in cell populations according to their replicative age after enrichment are shown in Supplementary Figure S7 After six hours of treatment, we observed that the culture became severely depleted of newborn cells and was dominated by mothers with 2 to 5 divisions.

The culture enriched in young mother cells and a control culture, where newborns were predominant, were synchronized at START with alpha-factor (no decreased sensitivity to alpha factor was detected in young mothers, Supplementary Figure S8) and allowed to progress into a new cell cycle. Samples were collected every 5 minutes and analyzed by flow cytometry. As observed in Figures 4B and 4C and Supplementary Figure S8, the young mother-enriched cultures (pink) progressed earlier into S phase compared to the newborn-enriched culture (blue). While the young mother-enriched culture exited G1 at 15-20 minutes, the exponential culture with a high proportion of newborn cells spent at least 5 more minutes before starting to enter S phase. This delay was consistent across five independent experiments and was statistically significant (Figure 4B). Interestingly, despite the earlier entry into S phase of young mother cells, both cultures exited G2/M phase at the same time, indicating that the S-G2-M phase of the cell cycle was longer in young mothers compared to newborn cells, at least in a substantial fraction of the cell population (Figures 4B and 4C).

As mentioned above, *WHI5* expression peaks during the S and G2/M phases. We hypothesized that the relatively longer S-G2-M duration during the first division cycles of those mother cells that exhibit this cell-cycle change could lead to higher Whi5 levels, due to the extended period in which *WHI5* is expressed. To test this, we carried out qPCR analyses of *WHI5* mRNA levels after alpha-factor synchronization and release. Specifically, we performed the same biotin-streptavidin purification and alpha-factor synchronization as described in the previous paragraph; however, this time, samples were collected at specific time points. To select these time points, we also analyzed samples from the previous experiment under a fluorescence microscope to determine the cell cycle stage of each cell more precisely by examining cell morphology and nuclei visualization (Figure 4D, right panel, and supplementary Table S2). We then chose the time points with the highest enrichment for each cell cycle phase category from four different replicates and analyzed them by RT-qPCR. As shown in Figure 4D (left panel), *WHI5* mRNA levels decreased from S phase to telophase in control cells but remained high in young mother cells, resulting in a highly significant accumulation in this latter category. This result indicates that young mother cells tend to accumulate Whi5 during the S-G2-M segment, before cytokinesis and the initiation of a new G1 phase of the cell cycle.

## Discussion

### Proliferative heterogeneity is coupled to mitotic division since first division rounds

Heterogeneity in clonal populations can take many forms, including very different proliferation rates^37^. We have focused on proliferative heterogeneity because, as previously described, microencapsulated isogenic single cells produced microcolonies with a wide range of sizes^14^. This size heterogeneity reflects the number of cells within the microcolony and, therefore, differences in the number of divisions undergone during its formation.

To understand proliferative heterogeneity, studies cannot be performed in culture populations but must analyze individual cells and colonies. We have developed a direct and powerful methodology to analyze the formation of individual microcolonies from a single founder cell by combining cell microencapsulation and confocal microscopy. Using this new experimental strategy, we have identified two different types of growth patterns associated to low proliferation rates. The most frequent pattern (observed in 83 out of 93 small microcolonies analyzed), which we call “isotropic” growth, is characterized by all cells proliferating at a low rate, with no particular cell in the microcolony exhibiting a stronger restriction on division. The second, less frequent pattern (observed in 10 out of 93 small microcolonies), which we call “anisotropic” growth, occurs when the founder cell undergoes multiple divisions, while its daughter cells either do not divide or experience severe restrictions on division. Anisotropic growth may be caused by mild stress experienced by founder cells during microencapsulation, as has been described in low-metal environments ^38^. The fact that only 10 out of 93 small microcolonies followed this growth pattern, while 83 out of 93 exhibited isotropic growth, suggests that the conclusions drawn in this study are not limited to stressed cell subpopulations.

Our analyses also provide relevant insights into the age of founder cells at the moment of encapsulation—that is, when slow-growing microcolonies were initiated—and its impact on the proliferative capacity of the future microcolony. According to our analyses, most slow-proliferating microcolonies were founded by non-newborn cells that had undergone fewer than five divisions, in both isotropic and anisotropic types. While the liquid cultures used for encapsulation exhibited the well-known age distribution of exponentially growing populations ^39–41^, characterized by a majority of newborn cells without bud scars (64%), followed by progressively smaller subpopulations of cells with one, two, three, or more bud scars (Figure 2E in red), the founder cells of slow-proliferating microcolonies did not follow this proportion. Instead, they were enriched in cells that had undergone one or more divisions before encapsulation (Figure 2E in grey). Moreover, the smaller the microcolony, the greater the number of division cycles previously undergone by the founder cells (Figure 2D). Finally, using the MEP system to enrich for non-newborn founder cells (one or two divisions), we observed a significant increase in slow-proliferating microcolonies (Figure 2F). We conclude that, once a newborn cell divides, there is significant probability that its descendant lineage will stably slow down its proliferation rate across generations. In other words, the restriction in proliferative capacity can be established at any stage of the mother cell’s lifespan, even during its first division cycle.

It is important to emphasize that the impact on proliferation rate that we describe occurs at the very early stages of the replicative lifespan. So far, this connection had been studied from the perspective of the final divisions that a mother cell undergoes before death, after more than 10 replicative cycles ^42–44^. Secondly, the impact of proliferation rate from the very first divisions provides a better understanding of the proliferative heterogeneity observed in exponential cultures (see our modelling results shown in Figure 3D), where the proportion of cells with more than ten divisions is quite insignificant. In fact, the restricted proliferation of young mother cells has a much greater impact on population fitness than functional deterioration at later ages and may play a substantial role in adaptation to changing environments^37^.

### Whi5 levels link proliferative heterogeneity to very early stages of the replicative lifespan

Cell cycle progression in eukaryotic cells requires complex regulatory networks to ensure functional success^45^. The timing of START in late G1 is regulated by the antagonistic interplay between the Cln3 G1 cyclin and the Whi5 inhibitor. It has been proposed that increased Cln3 synthesis, along with the dilution of Whi5 levels due to cell size expansion, controls START timing and cell size-dependent proliferation ^31^. As cell size increases, the Whi5 protein becomes progressively diluted. Once it reaches a certain threshold concentration—corresponding to an optimal division size—cells commit to division ^26^. In this work, we found a strong connection between Whi5 concentration and proliferation heterogeneity. By modulating its expression, we demonstrated a direct correlation between Whi5 levels and the proportion of slow-proliferating microcolonies (Figure 3B-C). Furthermore, the sorting results shown in Figure 3D-F indicate that higher Whi5 levels are associated with non-newborn cells that have undergone a small number of replicative cycles. These results confirm our previously published finding that the smallest microcolonies, mostly founded by non-newborn cells, overexpress Whi5^14^.

It has been previously described that Whi5 accumulates in very old aged cells, causing cell cycle delays and genomic instability that ultimately result in cell death^46^. However, this observation was based on mother cells that had undergone more than 10 division cycles, nearing the end of their lifespan. Here, we present results showing that young mother cells, after only 1 to 5 divisions, exhibit altered cell cycle phases duration in comparison to newborn cells (Figure 4A-C), and accumulate significantly higher levels of Whi5 (Figure 3F) due to increased expression prior to cytokinesis (Figure 4D). Our findings demonstrate that Whi5 accumulation can occur at any division of a mother cell, even at very early stages, and support the proposal of Whi5 as a key regulator of proliferation heterogeneity across generations.

Our work demonstrates a Whi5 dose-dependent regulation of proliferation rates, both at the moment of the establishment of a cell lineage and later on, across its following generations (Supplementary Figure S9A). However, is hard to understand this very strong effect of Whi5 concentrations on cell proliferation, only according to its so-far known function in coupling cell size and G1-S transition ^21, 25, 26, 31^. If it were the case, the impact of Whi5 concentration in proliferation would be rather limited (compare upper and middle panels in Supplementary Figure S9B). We think that, on top of this effect, increased Whi5 might repress the rate of cell growth, thereby provoking much more marked consequences in cell proliferation (Supplementary Figure S9B bottom panel). According to this view, reduced proliferation rate would result from a combination of reduced growth rate and increased size threshold to enter S phase.

It is difficult to imagine which molecular mechanism might be mediating this inhibitory role of Whi5 on growth rate. One possibility would be that Whi5, in addition of its specific repression of G1-S genes, caused by its SBF-mediated recruitment to them, might play a general repression of gene transcription across the genome. In this regard, our analysis of ChIP-seq data obtained by Swaffer et al.^47^ detected ubiquitous binding of Whi5 to gene bodies, in proportion to their transcription. This hypothetical general repressive role might be related to the sharp increase in specific cell growth rate that happens from G1 to S-G2-M coinciding with the migration of Whi5 out of the nucleus^48^. The increase in Whi5 levels might be due to stochastic noise in its expression, but it could also be linked to the timing of its expression during the cell cycle, which appears to be maximal in S/G2/M phases ^49^. Any delay in progression through S or G2/M could result in increased Whi5 levels. Our results show that newborn cells, after the first few division cycles (1–5 cycles), tend to extend the duration of this expression window and accumulate higher levels of *WHI5* mRNA before cytokinesis (Figure 4B-D).

It has recently been shown that Whi5 not only regulates the G1/S transition but also progression through the S/G2/M phases ^50^. Accordingly, alterations in the mitotic cycle during S/G2/M, such as those caused by the activation of checkpoint mechanisms, might lead to increased Whi5 levels. This, in turn, could induce subsequent cell cycle alterations in G1, potentially establishing a self-regulatory loop that propagates throughout the cell lineage. Such checkpoint activation has been described as playing a role in mitotic catastrophe during late aging ^51^. Our work suggests that this may be a more common phenomenon, which could be triggered at any stage of the replicative lifespan, explaining the high proliferative heterogeneity of cell populations.

The entire set of results presented in this work is consistent with and supports a model (Supplementary Figure S9A) in which a newborn cell with optimal proliferation capacity can undergo increased Whi5 expression at any division round, starting from the very first one, due to cell cycle alterations. This increase in Whi5 levels would negatively affect the proliferation of its progeny, favoring extensive heterogeneity across the population. We have previously shown that a reduced proliferation rate is linked to other phenotypic features, such as energy metabolism, since small microcolonies respire in the very same cultures where fast-growing cells are fermenting^14^. Slow growth is also associated with altered mRNA turnover and a different global gene expression pattern ^14^. In this sense, Whi5-dependent proliferation restrictions would not be merely a terminal phenomenon linked to aging and senescence, but rather a developmental process that allows cell differentiation in terms of growth rate within the population, thereby increasing its phenotypic diversity from the very first division cycle.

## Materials and Methods

### Strains

See supplementary table 3 for a list of all strains used in this work.

### Microencapsulation

Microencapsulation conditions were set to obtain single cell clonal cultures. First, cells were cultured to a O.D. _600nm_ of approximately 0.5 to ensure that it was at exponential growth phase. Microencapsulation of individual yeast cells in alginate microparticles was performed using a Cellena microencapsulator (Ingeniatrics) following the protocol already published ^14,15^. Cells were gently sonicated before microencapsulation to avoid aggregation. After microencapsulation, capsules were collected and cultured together in YPD for 13 hours at 30°C continuous stirring. Less than 1% of the capsules were seeded with more than one cell, producing more than one microcolony within the same particle. These capsules were discarded for further analyses.

### Mother cell enrichment by selection against newborns

DNY51 strain was used for mother cell enrichment as described ^28^. A graphical explanation of the Mother Enrichment Program developed by Lindstrom and Gottschling is shown in supplementary Figure S4A. DNY51 cells were grown at normal culture conditions to exponential phase, they were then collected and inoculated in 10ml of YPD (previously preheated at 30°C) with 1μM estradiol. They were cultured at 30°C for 3 hours. The estradiol was then washed out, and cells were resuspended in YPD to finally add them to the microencapsulation mixture. After microencapsulation, capsules were collected and cultured in YPD (without estradiol) for 13 hours at 30 °C.

### Modulation of Whi5 expression

KSY098-1 strain, kindly provided by Kurt Schmoller, allowed us to modulate *WHI5* expression, which in this strain is under the transcriptional control of an estrogen receptor derivative ^32^. A KSY089-1 culture was first growth to exponential phase (O.D._600nm_ of 0.5) and then microencapsulated following the usual protocol. After that, microcapsules were collected and then inoculated in flasks with YPD and estradiol at different concentrations (0, 2, 4, 6, 8 and 10 nM). The wildtype W303 control was separately encapsulated the same day. Finally, these and the W303 wildtype strain were cultured for 9,5 hours at 30° C at the indicated conditions.

### Light microscopy and microcolony size analysis

Microcolony size analyses were performed using a Leica DM750 optical microscope. Microcolonies were classified according to their size in small, medium and big microcolonies. As capsule diameter is stable (approximately 100μm), big microcolonies were considered those that surpass the radio of the capsule, medium microcolonies were those whose size range between half the radio and the radio, and those that were smaller than half the radio was considered small microcolonies (Supplementary Figure 1A).

### Calcofluor staining

For replicative age analysis bud scar staining was performed. First, cells were collected at exponential growth phase, fixed with 70% Ethanol and stored at 4°C for at least 2 hours. After that, cells were washed with PBS 1X and resuspended in a solution of PBS and calcofluor (Fluorescent Brightener 28, Sigma Aldrich) at a final concentration of 0.1 mg/ml. They were incubated for 5 to 10 minutes in dark conditions at room temperature and then washed again with PBS. Cells were finally resuspended in 100μl of PBS.

Microcolonies were also stained with calcofluor for small microcolonies founder cell identification. First, fixed microcolonies (YPD and Ethanol 70% 1:1) were filtrated and collected. They were washed with a TrisHCl-CaCl 2:1 solution and inoculated in 0.5 ml of calcofluor at 0.1 mg/ml concentration for 5 min at room temperature. After that, microcapsules were filtrated and washed again with TrisHCl-CaCl solution. Finally, they were collected and preserved in 0.5 ml of the same solution.

### Fluorescent microscopy and bud scar counting

Replicative age of the population was calculated by manually counting the number bud scars per cell using a direct fluorescent microscope Olympus BX-61. Bud scars of calcofluor-stained cells were detectable using the DAPI filter.

### Confocal microscopy and founder cells age analysis

Microcolonies stained with calcofluor were visualized in a A1R Confocal Nikon Microscope using the 60X oil objective and DAPI filter. Images of small microcolonies (n=187) were taken every 0.5 μM in Z-axis. Total number of cells and total number of bud scars were counted going through all Z axis images. Using the formula *number of budscars* − *number of cells* + 1 = *founder cell age* the replicative age of the small microcolonies founder cells at initial time (right before founding the small microcolony) was estimated. The resulting number was corrected when incoherent: very high numbers usually meant that the small microcolony was founded by two aged cells. These cases were taken out of the analysis. On the other hand, negative numbers probably meant that the number of bud scars was underestimated. Going through the genealogy of the small microcolonies, taken into consideration the number of buds observed on each cell, these numbers were corrected according to the most probable scenario (being usually the case of a newborn founder).

### Whi5-GFP cell sorting

Whi5-GFP strain was cultured to exponential growth in minimum complete media. Then, cells were collected at of OD_600nm_ of 0.5 and sorted in a BD FACSJazz™ cell sorter according to the GFP intensity detected. Cells with high Whi5 expression were considered the 5% with the highest GFP expression. After sorting, each sample was stained with calcofluor and analyzed for bud scar counting.

### Mother enrichment in synchronized cultures by biotin/streptavidin purification

Cells were collected at an O.D._600nm_ of 0.4. They were then centrifuged at 4,000 rpm for 4 minutes, washed twice with 1ml PBS (2 minutes at 6,000rpm) and incubated with 10 mg of biotin for 30 minutes. Half of the culture was collected as control cells (non-enriched in mother cells) and was incubated in YPAD for 30 minutes in agitation at 30°C. The rest of the culture was incubated for 6 hours in the same conditions (Supplementary Figure S8A). After each incubation, cells were treated with 1μg/ml of alfa-factor for 2 hours (or until the shmoo structure is visible under the microscope). After this, we performed a streptavidin extraction (New England Biolabs, 150 μl per sample) for 30 minutes at room temperature using streptavidin-linked magnetic beads. The supernatant was discarded and the cells attracted by the magnet were washed several times. After this step, cells were washed and inoculated into fresh YPAD media (previously heated at 30°C) with pronase at 50μg/ml. Samples were collected every 5 minutes for the following 90 minutes and fixed in 500 μl of 70% ethanol.

In these experiments, we ensured that mother cell enriched cultures had the mean replicative age of our interest by staining with calcofluor and counting the number of budscars.

### Analysis of cell size

Cell cultures were grown overnight to early log phase at 25°C. A 900 ml sample of each culture was fixed with 100 ml of 37% formaldehyde for 30 min and then washed twice with PBS containing 0.02% Tween-20. Cell size was measured using a Coulter Counter (BECKMAN COULTER MULTISIZER 3, Beckman Coulter) as previously described (Jorgensen et al., 2002). In brief, cells were diluted into 10 ml diluent (0.9% NaCl) and sonicated for 3 s before cell sizing. Each plot is the average of three independent biological replicates in which three independent technical replicates were averaged.

### Propidium iodide staining and flow cytometry

Fixed cells were washed twice with 1ml of sodium citrate 50mM and then treated with 500μl of sodium citrate with RNAase overnight. After this, 500 μl of a solution of propidium iodide (PI)(4μg/ml) and sodium citrate 50 nM was added to every sample. Cells were incubated for 30 min at room temperature and dark conditions.

For flow cytometry, samples were sonicated during 3 cycles of 1 min (30 seconds of sonication for every cycle). After this, cell cycle analysis was performed using a BD FACSCanto™ Clinical Flow Cytometry System. Cells were classified according to their PI content, which correlates with DNA content. This allows sorting cells into G1 cells (1 content of DNA) and G2/M (2 contents of DNA).

### *WHI5* RTqPCR analysis in synchronized young mother cells

In order to determine Whi5 mRNA levels for every cell cycle phase, samples of synchronized cultures were collected and then analyzed by RTqPCR. First, cultures were collected at exponential growth and enriched in mother cells using the same protocol described above. After alfa-factor treatment, cells were enriched in mother cells (early aged cells) through streptavidin extraction and then inoculated into YPAD and pronase. 15 ml samples were collected at the indicated times for every experiment and then washed and frozen in liquid nitrogen. For every timepoint, 20 ml samples were used for RT-qPCR and an additional sample was analyzed by flow cytometry and microscopy, in order to check the actual cell cycle phase.

Quantitation of *WHI5* mRNA levels by RT-qPCR was carried out using the following primers: 5’ GGCTGCGCGTCGGTCTAC 3’ and 5’ TGCAGCTTGACTAACGCGTAAT 3’. It was performed in a LightCycler® 480 Instrument II thermocycler (Roche). For each reaction we used 4 μl of sample and 6 μl of a mixture with forward and reverse oligonucleotides and Sybr Green Premix.

### Mathematical modelling of growth rates

To analyze the bulk growth rates of KSY098 cultures upon Whi5 upregulation, we measured OD vs. time in exponential liquid culture without estradiol, and with 2, 4, 6, 8 and 10 nM estradiol (three biological replicates per concentration). We performed a linear regression between the Log base 2 of OD and time, and the average cell doubling time is estimated as the inverse of the slope of this linear regression (Supplementary Figure S5D).

To estimate the bulk growth rates from the semi-qualitative data on colony size distribution (the number of small, medium and big microcolonies) in Fig. 3C across estradiol concentrations, we assumed that encapsulated small, medium and big microcolonies had an average of 23, 343 and 1172 cells, respectively, after 13 hours of growth starting from a single cell. Then, for a given estradiol concentration, the average cell doubling time in minutes was obtained using the formula:

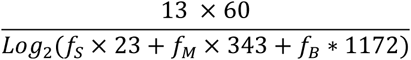

where *f_s_, f_M_, f_B_* is the measured average fraction of small, medium and big microcolonies in Fig. 3C, respectively.

## Supporting information

Supplementary figures_Delgado-Roman

## Author’s Contributions

I.D.R. performed most experiments, analyzed data, contributed to discussions and prepared the figures; M.J.G.M. performed specific experiments at the final stages and contributed to figures; L.D.R. performed research at the initial stages; M.C.M.C. and S.C. designed research and discussed results; M.C.M.C. wrote the manuscript.

## Acknowledgments

We thank all the people of the IBiS Gene Expression lab for helpful discussion and all technicians from IBiS and CITIUS facilities. We are grateful to Rafael Lucena for his valuable discussions and support with the cell size analysis. We thank María Siles Mudarra for her help with the estradiol cytometry experiments. We deeply thank Kurt Schmoller and Michael Chang and Daniel Gottschling for KSY098-1 and DNY51 strains, respectively. We also thank our colleagues of IBiS and the Genetics Department of Universidad de Sevilla for their support.

