## Supplementary figures_Delgado-Roman for "Mother cells can establish slow growing lineages in clonal populations since their earliest division cycles"

| Colony ID. | Number of cells | Number of bud scars | Founder cell age | Colony ID. | Number of cells | Number of bud scars | Founder cell age |
|---|---|---|---|---|---|---|---|
| 1 | 13 | 15 | 3 | 49 | 27 | 26 | 0 |
| 2 | 44 | 50 | 7 | 50 | 29 | 28 | 0 |
| 3 | 6 | 6 | 1 | 51 | 11 | 13 | 3 |
| 4 | 16 | 15 | 0 | 52 | 26 | 27 | 2 |
| 5 | 38 | 37 | 0 | 53 | 25 | 24 | 0 |
| 6 | 28 | 31 | 4 | 54 | 34 | 34 | 1 |
| 7 | 22 | 22 | 1 | 55 | 36 | 37 | 2 |
| 8 | 68 | 67 | 0 | 56 | 26 | 25 | 0 |
| 9 | 5 | 8 | 4 | 57 | 28 | 27 | 0 |
| 10 | 15 | 16 | 2 | 58 | 10 | 9 | 0 |
| 11 | 69 | 68 | 0 | 59 | 17 | 17 | 1 |
| 12 | 24 | 27 | 4 | 60 | 26 | 25 | 0 |
| 13 | 9 | 12 | 4 | 61 | 25 | 26 | 2 |
| 14 | 25 | 24 | 0 | 62 | 17 | 18 | 2 |
| 15 | 38 | 37 | 0 | 63 | 9 | 10 | 2 |
| 16 | 62 | 61 | 0 | 64 | 41 | 42 | 2 |
| 17 | 15 | 14 | 0 | 65 | 27 | 28 | 2 |
| 18 | 13 | 14 | 2 | 66 | 24 | 23 | 0 |
| 19 | 10 | 9 | 0 | 67 | 25 | 26 | 2 |
| 20 | 23 | 22 | 0 | 68 | 15 | 15 | 1 |
| 21 | 44 | 43 | 0 | 69 | 9 | 9 | 1 |
| 22 | 34 | 33 | 0 | 70 | 11 | 13 | 3 |
| 23 | 21 | 20 | 0 | 71 | 9 | 8 | 0 |
| 24 | 21 | 22 | 2 | 72 | 10 | 11 | 2 |
| 25 | 12 | 13 | 2 | 73 | 6 | 5 | 0 |
| 26 | 50 | 50 | 1 | 74 | 24 | 25 | 2 |
| 27 | 21 | 20 | 0 | 75 | 26 | 27 | 2 |
| 28 | 33 | 32 | 0 | 76 | 15 | 14 | 0 |
| 29 | 26 | 26 | 1 | 77 | 6 | 6 | 1 |
| 30 | 46 | 45 | 0 | 78 | 8 | 7 | 0 |
| 31 | 30 | 29 | 0 | 79 | 26 | 25 | 0 |
| 32 | 11 | 12 | 2 | 80 | 14 | 15 | 2 |
| 33 | 11 | 11 | 1 | 81 | 7 | 8 | 2 |
| 34 | 12 | 15 | 4 | 82 | 6 | 8 | 3 |
| 35 | 12 | 16 | 5 | 83 | 10 | 9 | 0 |
| 36 | 45 | 46 | 2 | 84 | 5 | 6 | 2 |
| 37 | 8 | 9 | 2 | 85 | 17 | 16 | 0 |
| 38 | 36 | 35 | 0 | 86 | 18 | 17 | 0 |
| 39 | 38 | 37 | 0 | 87 | 9 | 9 | 1 |
| 40 | 22 | 21 | 0 | 88 | 22 | 21 | 0 |
| 41 | 18 | 17 | 0 | 89 | 10 | 9 | 0 |
| 42 | 40 | 41 | 2 | 90 | 6 | 6 | 1 |
| 43 | 48 | 49 | 2 | 91 | 9 | 8 | 0 |
| 44 | 20 | 20 | 1 | 92 | 29 | 30 | 2 |
| 45 | 28 | 27 | 0 | 93 | 5 | 6 | 2 |
| 46 | 43 | 44 | 2 | 94 | 24 | 24 | 1 |
| 47 | 34 | 34 | 1 | 95 | 17 | 16 | 0 |
| 48 | 28 | 31 | 4 | 96 | 7 | 10 | 4 |

**Table S1. Quantitative analysis of slow-proliferating microcolonies.** For each microcolony, the number of cells, total number of bud scars counted, and the calculated age of the founder cell (at the time of founding) are provided. The replicative age of each founder cell was estimated according to the formula: number of bud scars - number of cells + 1 = Founder cell age. All images used in this study can be found at http://www.ebi.ac.uk/biostudies/ with the accession number S-BSST1071.

| | | Control | | | | | Mother-enriched | | | | |
|---|---|---|---|---|---|---|---|---|---|---|---|
| **Cell cycle phase** | | 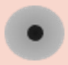 | 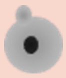 | 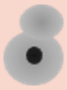 | 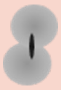 | 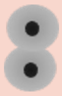 | 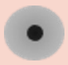 | 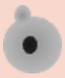 | 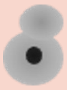 | 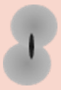 | 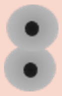 |
| **START (t0 min)** | **R1** | 70 | 29 | 1 | 0 | 0 | 78 | 12 | 10 | 0 | 0 |
| | **R2** | 83 | 14 | 3 | 0 | 0 | 83 | 4 | 13 | 0 | 0 |
| | **R3** | 90 | 15 | 0 | 0 | 0 | 83 | 9 | 8 | 0 | 0 |
| **S (t20/25 min)** | **R1** | 33 | 49 | 18 | 0 | 0 | 34 | 43 | 23 | 0 | 0 |
| | **R2** | 50 | 42 | 8 | 0 | 0 | 33 | 55 | 12 | 0 | 0 |
| | **R3** | 30 | 69 | 1 | 0 | 0 | 43 | 42 | 15 | 0 | 0 |
| **G2/M (t40/45 min)** | **R1** | 25 | 7 | 60 | 6 | 2 | 18 | 5 | 50 | 11 | 16 |
| | **R2** | 20 | 11 | 58 | 11 | 0 | 17 | 6 | 59 | 9 | 9 |
| | **R3** | 26 | 5 | 60 | 9 | 0 | 27 | 6 | 55 | 7 | 5 |
| **Telophase (t60/65 min)** | **R1** | 10 | 6 | 22 | 14 | 48 | 17 | 3 | 25 | 5 | 50 |
| | **R2** | 11 | 1 | 18 | 13 | 57 | 21 | 1 | 22 | 8 | 48 |
| | **R3** | 20 | 1 | 30 | 11 | 38 | 28 | 1 | 30 | 5 | 36 |

**Table S2: Cell-cycle distribution of control and young mother-enriched cells at different time points after release from α-factor synchronization.** Three replicates of cells, prepared as described in Figure 4B, were sampled for analysis after staining with propidium iodide and manual counting under a fluorescence microscope. Time points were selected for each replicate to align with START, S phase, G2/M and telophase. The cells obtained from these time points were then used for RT-PCR analysis of *WHI5* mRNA, as described in Figure 4D.

| Name | Genotype | Source |
|---|---|---|
| **BY4741** | *MATa; his3Δ1; leu2Δ0; met15Δ0; ura3Δ0* | Euroscarf |
| **DNY51** | *MATa can1::PSTE2-Sp_his5 leu2Δyp1Δ met15Δ hoΔ::PSCWII-cre-EBD78-NATMX loxP-ubc9-LOXp-Leu2 loxP-CDC20-Intron-loxP-HPHMX* | From Lindstrom & Gottschling, 2009 |
| **KSY098-1** | *MATa; his3::LexA-ER-AD-TF-HIS3 whi5::kanMX6-LexApr-Whi5-mCherry-ADH1term-LEU2* | Prof. Schmoller lab, Helmholtz Zentrum München, Germany |
| **W303** | *MATa leu2Δ3 112 trp1-1 can1-100 ura3Δ1 ade2Δ1 his3Δ11 15* | Lab collection |
| **Whi5-GFP** | *MATa his3Δ1 leu2Δ0 met15Δ0 ura3Δ0 Whi5-GFP::HIS3* | Thermo Fisher Scientific GFP yeast clone collection |

**Table S3. Strains used in this work**

Figure S1

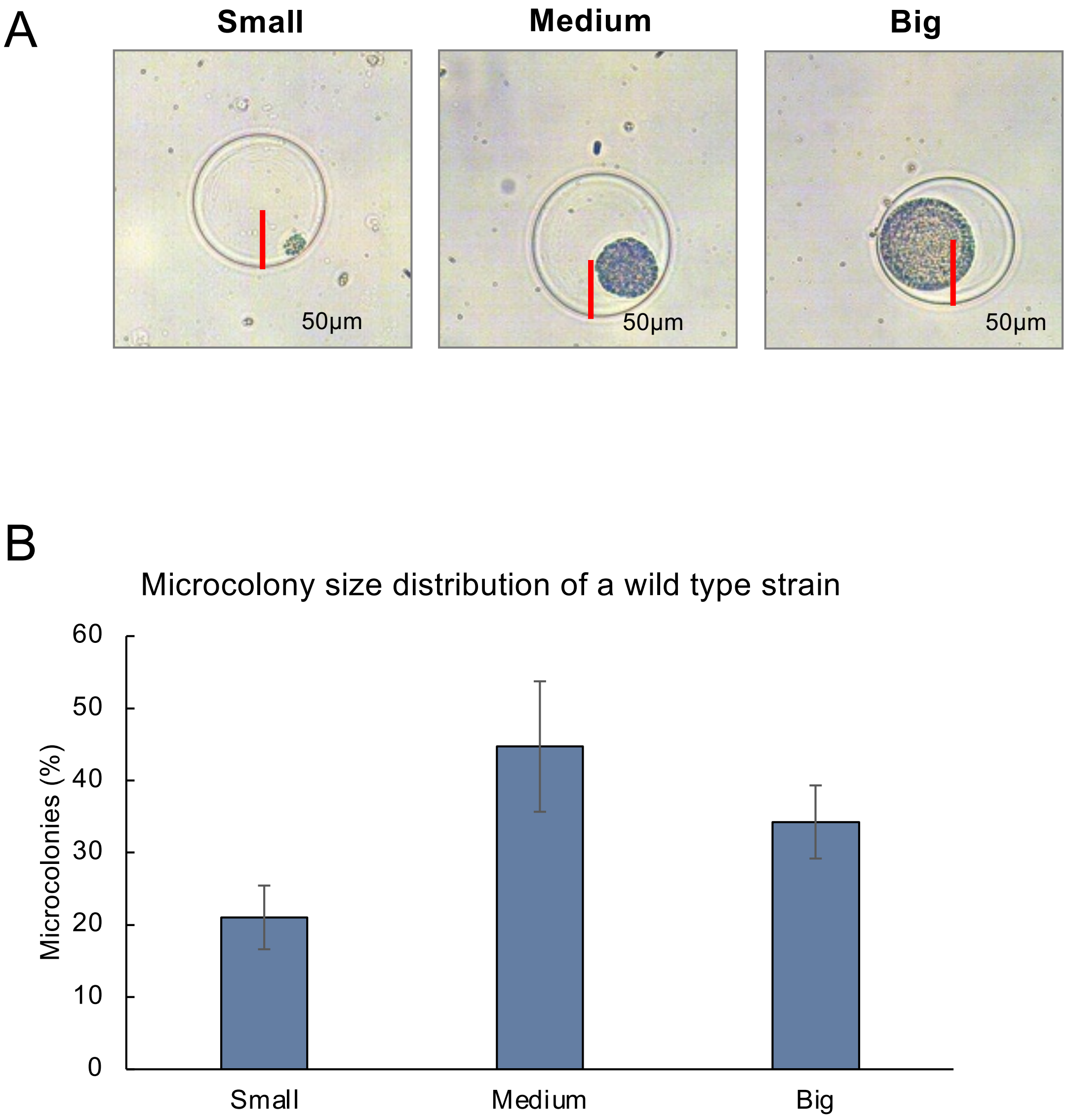


**Figure S1: Semi-quantitative analysis of microcolony size distribution.** (A) Guide diagram for microcolony size classification. Small microcolonies are smaller than half the capsule radius. Large microcolonies exceed the length of the capsule radius. Medium microcolonies range from half to the full capsule radius in length. (B) An example of a microcolony size distribution in a wild-type strain.

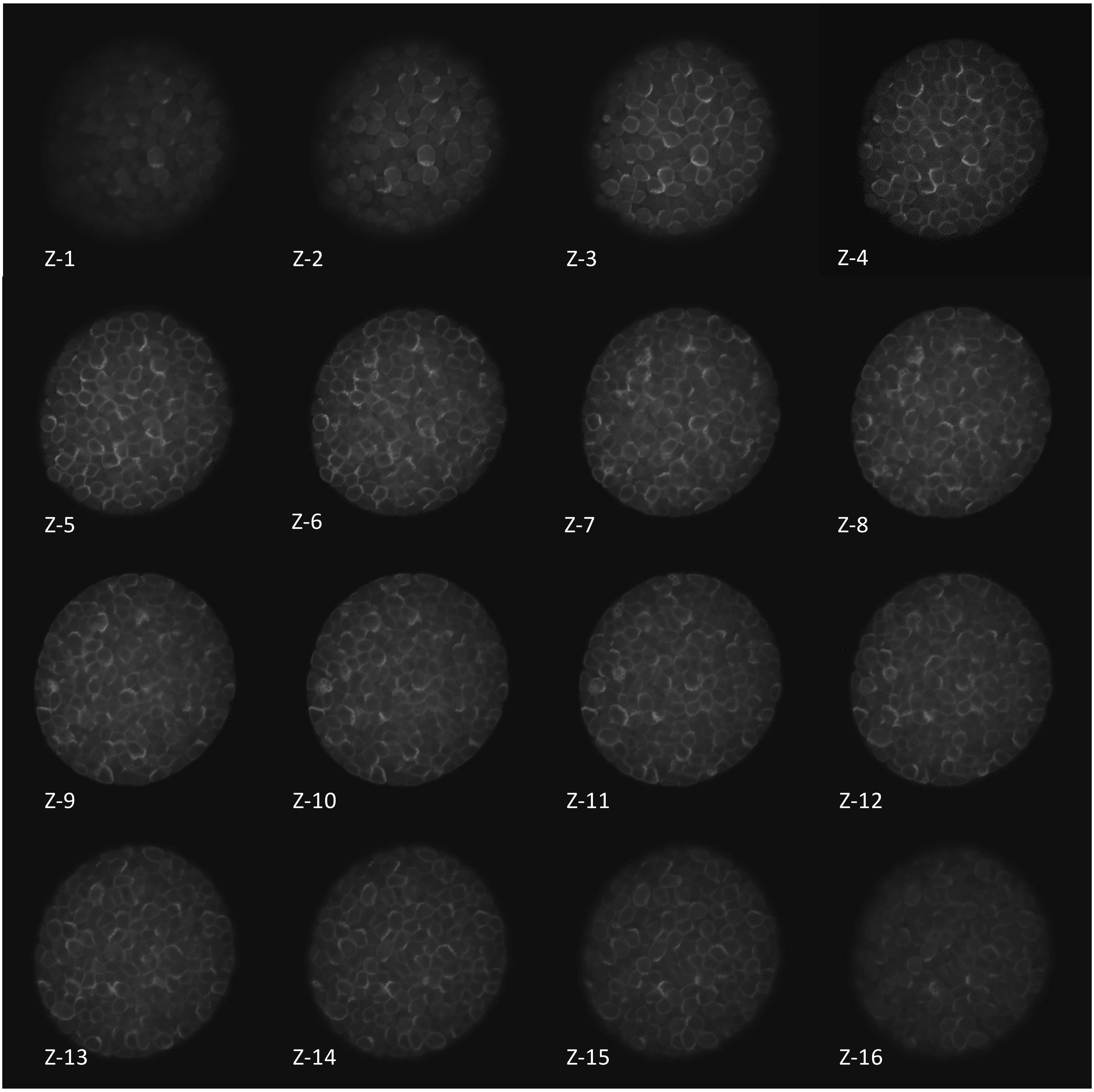


**Figure S2: Confocal microscopy images of a big microcolony stained with calcofluor.** Images were taken every 0.5 micrometre intervals along on Z axis.

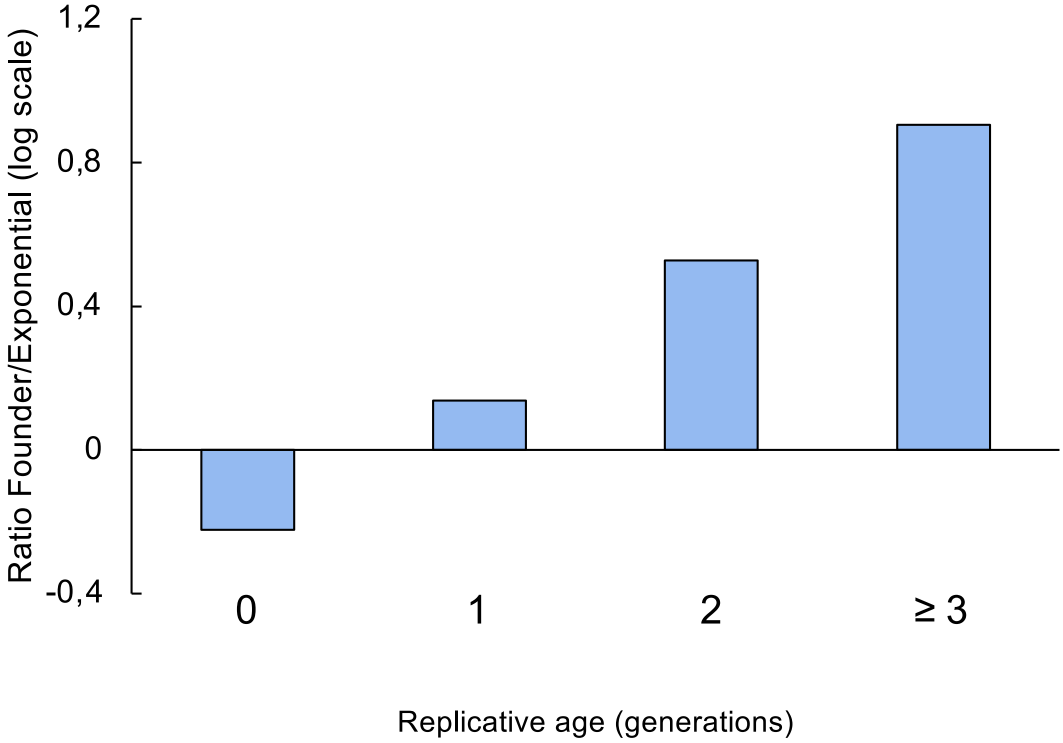


**Figure S3: Slow-proliferating microcolonies are founded more frequently founded by young mother cells than would be expected based on the age distribution of cells in an exponential culture.** This figures shows the ratio of small microcolony founder cells with replicative ages of 0, 1, 2 or 3 or more to the proportion of cells in an exponential culture with the same replicative age.

Figure S4

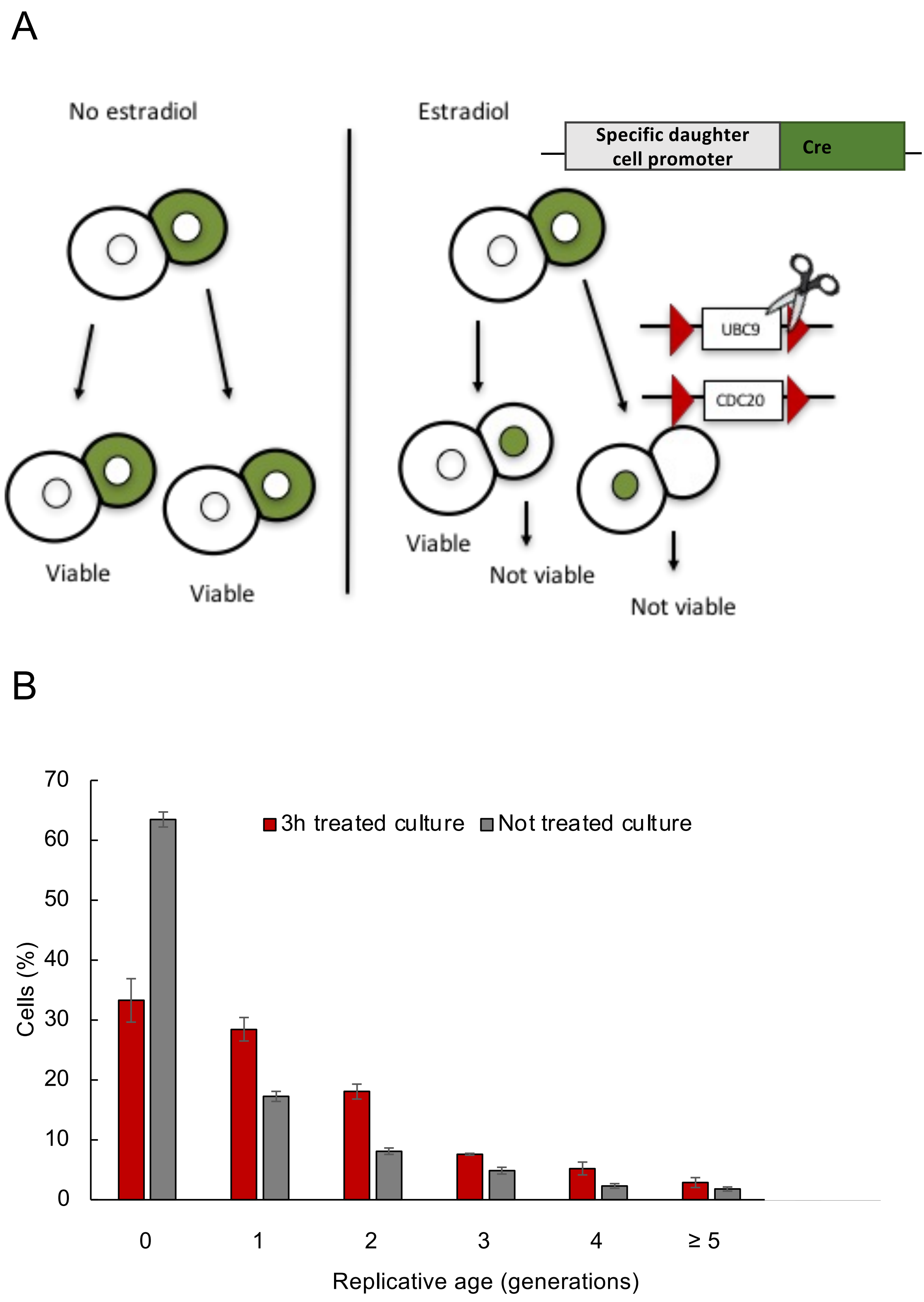


**Figure S4: Follow-up of the enrichment of a culture of young mother cells after the Mother Enrichment Programme (MEP).** (A) Graphical explanation of the MEP. A yeast strain with Cre recombinase under the control of a specific daughter cell promoter. In the presence of estradiol, the Cre recombinase enters the daughter cell nucleus and deletes two genes essential for the cell cycle. This results in the daughter cells becoming irreversibly arrested, while the mother cells continue to grow unaffected (based on Lindström & Gottschling, 2009). (B) DNY51 cells were treated with estradiol for three hours. The resulting cells (red) were counted for bud scars after calcofluor staining. A non-treated control is shown in grey.

Figure S5

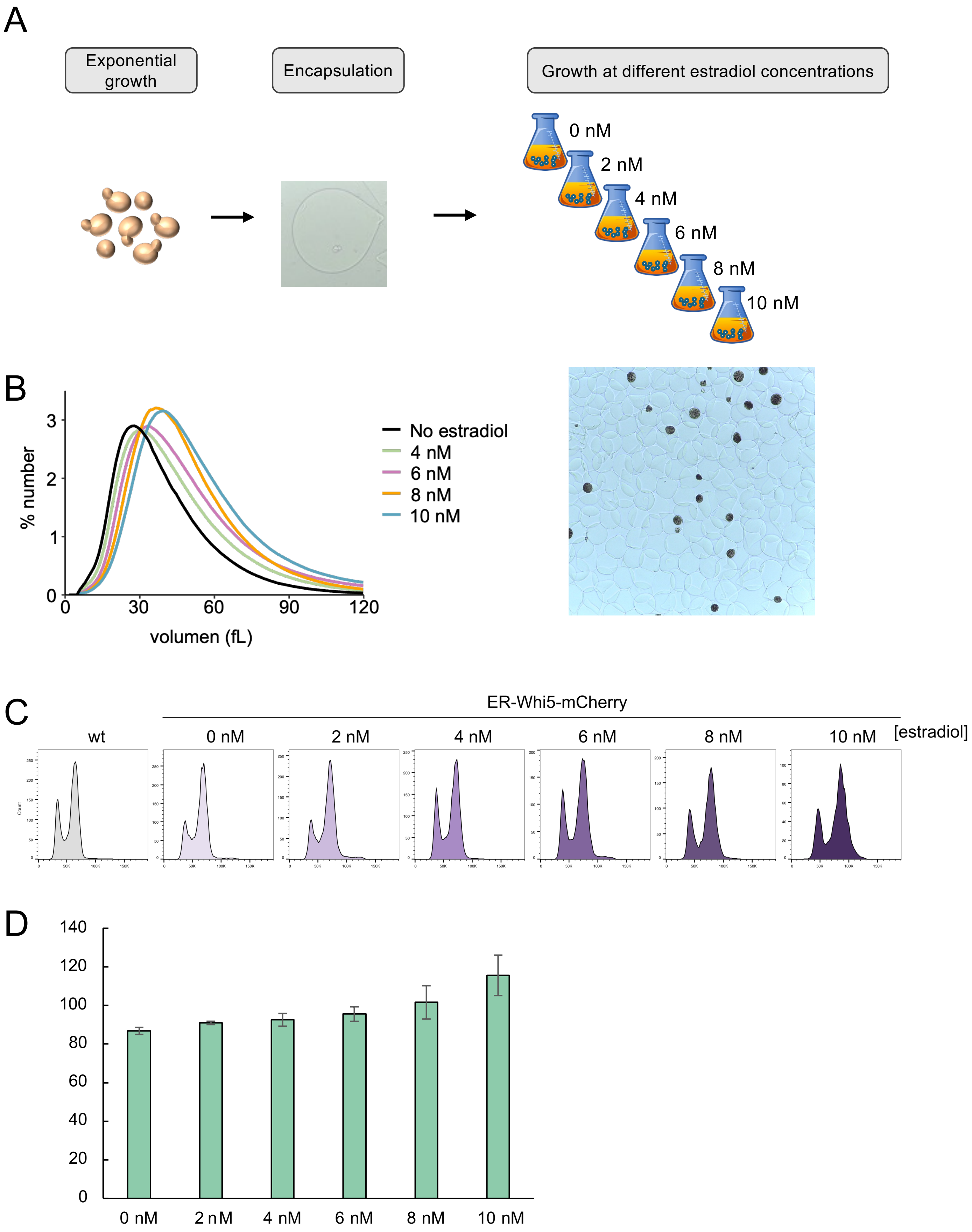


**Figure S5: Modulation of Whi5 levels in encapsulated cells.** (A) Experimental procedure for obtaining capsule cultures with increasing Whi5 concentrations using the KSY098 strain with various concentrations of estradiol (2 nM, 4 nM, 6 nM, 8 nM or 10 nM), or YPD medium without estradiol. The cultures were then incubated under standard growth conditions to allow microcolony proliferation. (B) Left panel shows cell volume distributions after growth at different estradiol concentrations. Right panel shows microcolonies of different sizes in an exponentially growing culture (C) Cell cycle profiles of cells grown under different estradiol concentrations. (D) Average cell-doubling time in mins for 0, 2, 4, 6, 8,10 nM estradiol in liquid culture with errors bars denoting 95% confidence intervals.

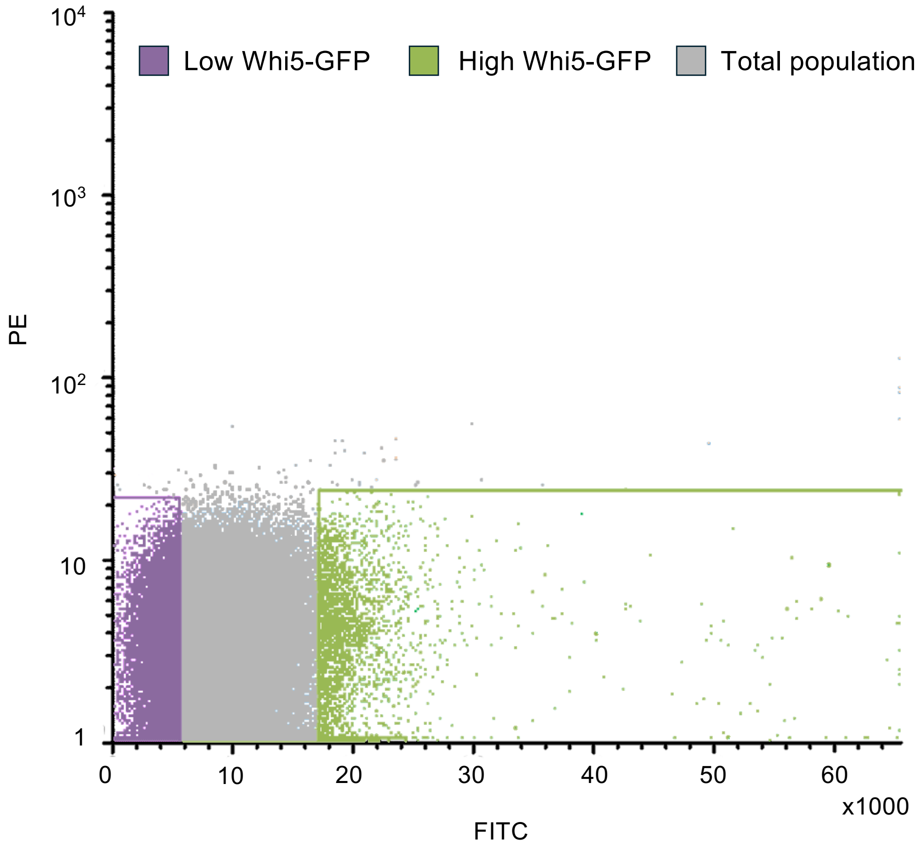


**Figure S6: Cell sorting of a Whi5-GFP strain based on GFP intensity.** Cell sorting based on the Whi5-GFP signal. Purple represents cells sorted into the 'low Whi5 expression' group, and green represents cells sorted into the 'high Whi5 expression' group.

Figure S7

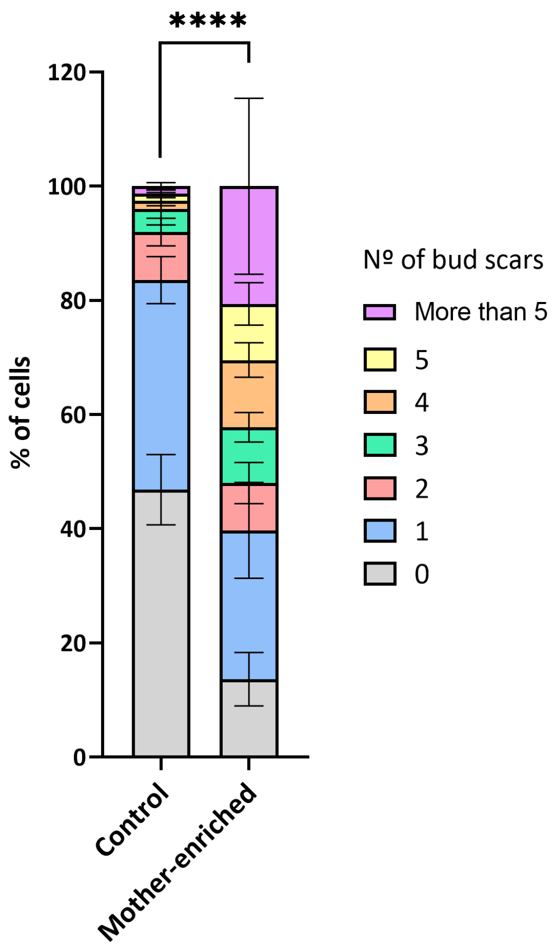


**Figure S7: Replicative age distribution in a 6-hour mother-enriched culture using the biotin-streptavidin method.** Compared to a control culture, the mother-enriched culture contained fewer than one-third of the number of newborn cells, consisting mostly of cells that had undergone 2 to 5 divisions.

Figure S8

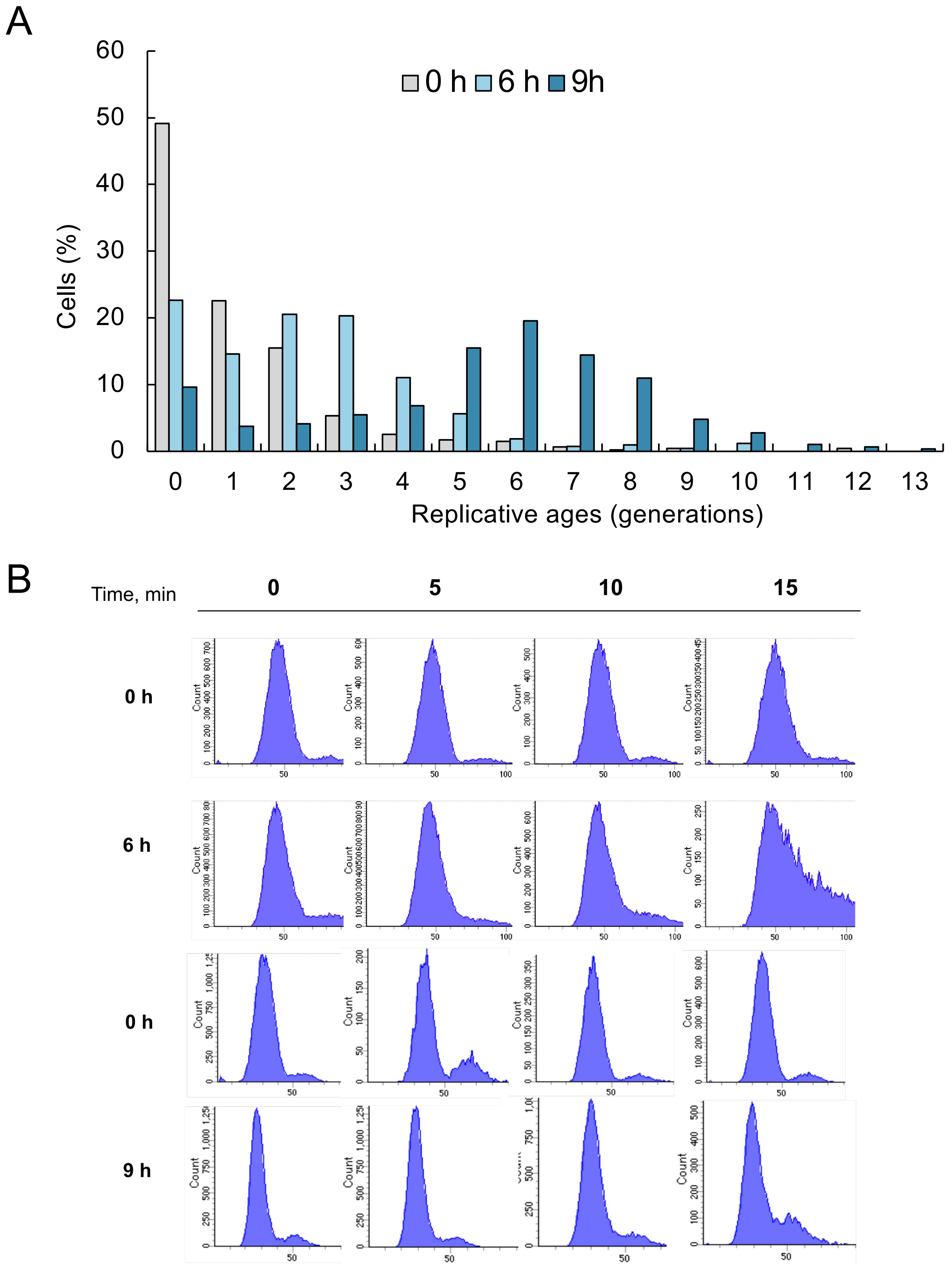


**Figure S8: Analysis of synchronized cultures enriched for early-aged cells.** (A) Distribution of cells according to their replicative age in different mother-enriched cultures. (B) Flow cytometry profiles, comparing cultures enriched for 6 or 9 hours with a non-enriched control (0 h) after α-factor release.

Figure S9

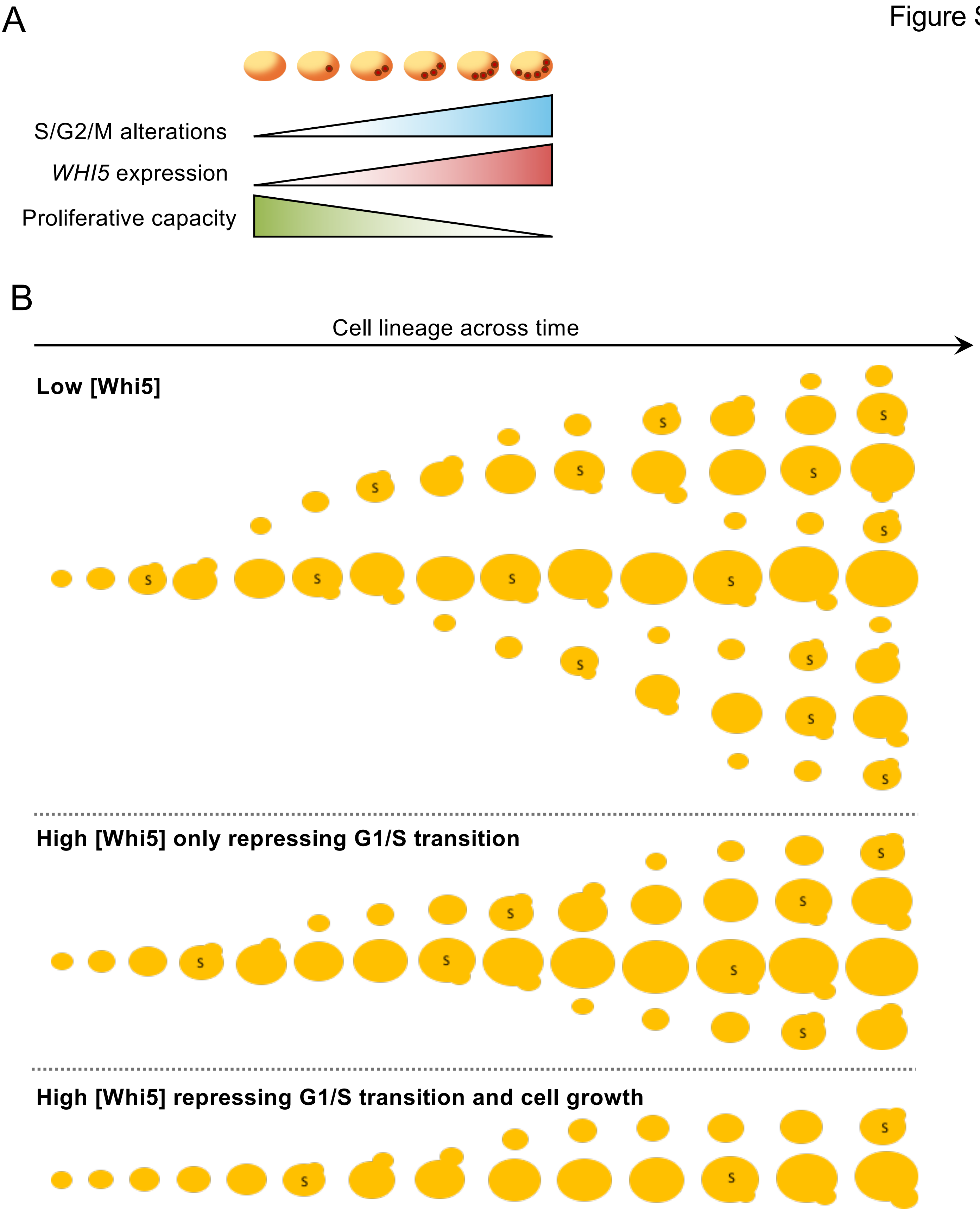


**Figure S9: Influence of stochastic Whi5 expression on proliferation the model.** (A) With each division of the young mother cell, the probability of an increased Whi5 concentration rises. In our model, this increased Whi5 concentration extends the generation time of the cell and is transmitted to the daughter cell, thereby triggering a positive feedback loop of Whi5 expression and slow growth across the cell lineage. (B) Differential proliferative capacities of mother cells and their progeny depending on their initial Whi5 levels, and its repressive role on G1/S transition only, or also on cell growth.
